# Transgenic mouse models for investigating human *DUX4* expression during development and its roles in FSHD pathophysiology

**DOI:** 10.1101/2025.08.22.671867

**Authors:** Yosuke Hiramuki, Charis L. Himeda, Peter L. Jones, Takako I. Jones

## Abstract

Facioscapulohumeral muscular dystrophy (FSHD) is an autosomal dominant myopathy caused by aberrant expression of the *DUX4* retrogene, and it affects skeletal muscles primarily in the face, shoulder, and limbs. In healthy individuals, *DUX4* is expressed in early development and is subsequently silenced in most somatic tissues. The spatiotemporal pattern of DUX4 mis-expression beyond the cleavage stage in FSHD is poorly understood because *DUX4* is not well conserved beyond primates. Here, we generated Cre reporter mouse lines with human *DUX4* regulatory elements to investigate the cell lineages derived from *DUX4*-expressing cells in embryos and adults. Intriguingly, we found that *DUX4*-expressing cell lineages were present in embryonic forelimb, hindlimb, and face. In adults, the reporter was expressed strongly in testis and to a lesser extent in other tissues, including weak, sporadic expression in skeletal muscles, reminiscent of mosaic DUX4 expression in FSHD. Within skeletal muscles, DUX4 lineage cells include pericytes, an interstitial cell that contributes to muscle regeneration and repair. Overall, this study introduces a new research tool for the field, and provides new insight into potential developmental mechanisms underlying FSHD pathophysiology.

**Summary statement:** Cre reporter mouse lines with human *DUX4* regulatory elements are discovery tools for developmental processes and mechanisms underlying FSHD pathophysiology.

## Introduction

Facioscapulohumeral muscular dystrophy (FSHD) is the third most common autosomal dominant muscle disorder with a prevalence of ∼1:8,500 – 15,000 (Deenen et al., 2014; Deenen et al., 2025; Orphanet, 2024). While FSHD pathology is highly variable, it typically affects muscles of the face, shoulder blades, and upper arms initially, then progresses to the abdomen and lower legs, with much of the weakness and atrophy appearing asymmetrically. Ultimately, all skeletal muscles are at risk of eventually becoming affected (Mul, 2022; Padberg, 1982). In addition, extra-muscular disease manifestations can occur in the severe infantile form of FSHD, including high frequency hearing loss and retinal abnormalities (Chen et al., 2020; Goselink et al., 2017; Klinge et al., 2006). Many FSHD clinical characteristics are unusual among neuromuscular diseases, and the underlying pathological mechanism(s) are not fully understood.

All forms of FSHD are caused by the loss of local epigenetic repression resulting in aberrantly increased expression of the *DUX4* (double homeobox 4) retrogene from the chromosome 4q35 D4Z4 array (Himeda and Jones, 2019; Lemmers et al., 2010; Snider et al., 2010). *DUX4* encodes a DUXC family double homeodomain transcription factor that, similar to other *DUXC* genes, initiates an early zygotic gene expression program in cleavage stage embryos, after which it is silenced in healthy somatic cells (De Iaco et al., 2017; Dixit et al., 2007; Gabriels et al., 1999; Hendrickson et al., 2017; Kowaljow et al., 2007; Leidenroth et al., 2012; Nip et al., 2023; Whiddon et al., 2017). However, in FSHD, *DUX4* is epigenetically de-repressed, leading to aberrant upregulation of its mRNA, protein, and target genes in skeletal muscles. Interestingly, aberrantly increased expression of DUX4 target genes has been detected in human FSHD1 fetal muscle biopsies as early as 14 weeks (Ferreboeuf et al., 2014). Initial expression of DUX4 establishes an epigenetic signature at its target genes that primes them for expression upon later exposure to DUX4 (Resnick et al., 2019); this suggests that there could be a developmental role for early DUX4 expression in dictating later FSHD pathology. The only healthy adult tissue where DUX4 protein is known to be expressed is testis, although *DUX4* mRNA has been reported in the thymus, cultured human keratinocytes (Gannon et al., 2016), and in vitro derived osteoblasts (de la Kethulle de Ryhove et al., 2015). Overall, very little is known about the pattern of *DUX4* expression in the healthy or FSHD states in vivo. Although all placental organisms have a *DUXC* ortholog, human *DUX4* is primate-specific, rendering it difficult to study developmentally in traditional model organisms, and all FSHD animal models are engineered transgenics (Bosnakovski et al., 2017; Bosnakovski et al., 2022; Jones and Jones, 2018; Jones et al., 2016; Pakula et al., 2019; Wuebbles et al., 2010). Thus, any developmental role for endogenous DUX4 expression in mediating typical or atypical FSHD pathology is still unknown.

Here we took a transgenic approach to investigate the developmental expression of *DUX4.* Previously, we characterized two *DUX4* myogenic enhancer (DME) regions centromeric to the 4q35 D4Z4 array that are required for *DUX4* expression in myogenic cells (Himeda et al., 2014). In addition, others have identified putative regulatory elements within the D4Z4 repeat itself (Gabriels et al., 1999; Ottaviani et al., 2010). Using these findings, we assembled a reporter transgene containing cis human *DUX4* regulatory elements including the DME regions and replaced the *DUX4* coding sequence with cre recombinase/EGFP to eliminate the cytotoxic effect of DUX4 and allow cell lineage tracing during embryogenesis and in adult tissues with desired floxed reporter lines. Using the sensitive floxed lacZ reporter mouse, we observed positive cell lineages in face and limbs of embryos, and in testis, heart, and skeletal muscles of adults, and identified blood vessel-associated pericytes as a previously unreported DUX4-expressing cell lineage.

## Results

### Generation of pJ2-Cre:EGFP mice

We are interested in understanding the developmental expression of DUX4 in FSHD; however, due to lack of evolutionary conservation of orthologs outside of primates, the synteny of the region, and genome organization of the repeat array, there were no suitable in vivo vertebrate developmental systems available. Although the *DUX4* coding sequence itself is primate-specific, the *DUX* family of double homeodomain protein encoding genes as well as D4Z4-like repeats are conserved between primates and mice (Clapp et al., 2007; Leidenroth et al., 2012; Leidenroth and Hewitt, 2010), suggesting that mice may have the capacity to appropriately regulate the expression of human *DUX4*. Therefore, we created a transgene consisting of the known endogenous *cis* transcriptional *DUX4* regulatory elements to generate transgenic mice that recapitulate developmental *DUX4* expression profiles. The transgene construct, pJ2-Cre:EGFP, consists of the two *DUX4* myogenic enhancers (1230-bp DME1 and 2100-bp DME2) (Himeda et al., 2014), a single D4Z4 repeat unit (RU) containing the core *DUX4* promoter elements and any regulatory elements within the repeat (Gabriels et al., 1999; Ottaviani et al., 2010), the CreEGFP fusion gene in place of the *DUX4* open reading frame, and a β-globin PAS (Fig. 1A). Specifically, the transgene contains the endogenous 5700-bp sequence proximal to the most centromeric 4q35 D4Z4 RU, from the KpnI site to DME1, of human chromosome 4q35. DME2, located ∼19 kb proximal to the most centromeric 4q35 D4Z4 RU, was placed upstream and directly adjacent to DME1.

**Figure 1.**
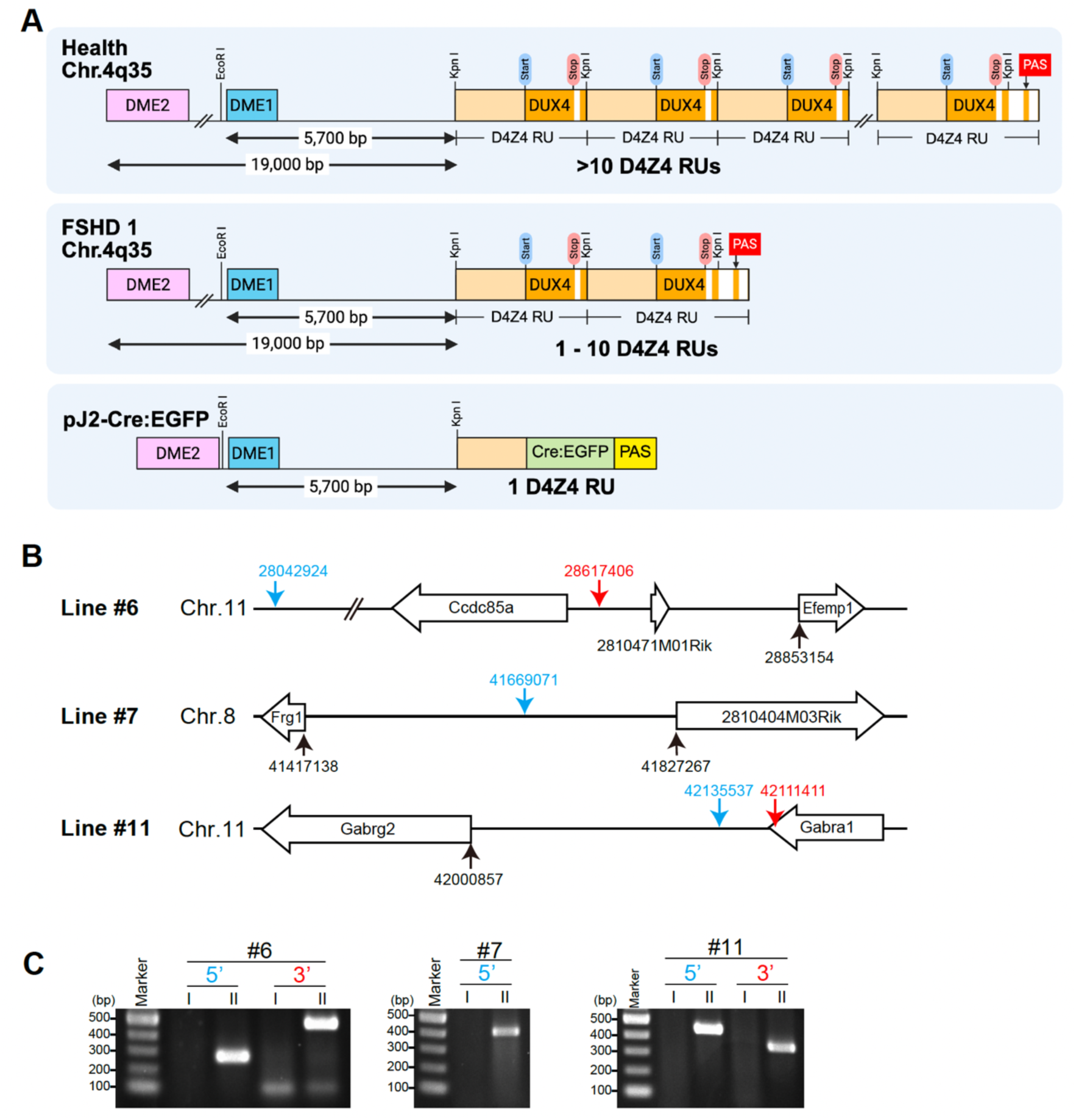
Generation of pJ2-Cre:EGFP mice. A) Diagram of the human chromosome 4q35 *D4Z4/DUX4* locus in healthy, FSHD1, and the pJ2-Cre:EGFP construct. Each D4Z4 RU (3303 bp) contains the *DUX4* exon 1 and exon 2 (orange), and the last D4Z4 RU has an exon 3 (orange) that includes the PAS used in FSHD. A 1230 bp DME1 (blue), a 2100 bp DME2 (pink), and D4Z4 without *DUX4* (light orange) are indicated. For pJ2-Cre:EGFP, *DUX4* is replaced by a Cre:EGFP fusion gene (green) and a β-globin PAS (yellow) B) Diagrams of integration sites of the three lines. Blue and red arrows indicate the 5’ and 3’ integration sites of transgenes, respectively. Genomic locus position was compared to the mouse mm10 sequence. C) PCRs for validating the transgene integration sites. I and II indicate *R26^NZG/+^* and *pJ2-Cre:EGFP; R26^NZG/+^*mice, respectively, for lines #6, #7, and #11. For line #6, the 5’ site (283 bp) and 3’ site (495 bp) products are shown; for line #7, only the 5’ site (411 bp) was identified. For line #11, the 5’ (453 bp) and 3’ (365 bp) products are shown. DME: <u>D</u>UX4 <u>m</u>yogenic <u>e</u>nhancer.

Transgenic mice were generated by random integration of the linearized construct into B6;SJL hybrid eggs using standard protocols. The resultant progeny were backcrossed to C57BL/6J and screened for the transgene by PCR. During backcrossing, three independent lines of pJ2-Cre:EGFP mice (#6, #7, and #11) were selected for relatively normal Mendelian inheritance of the transgene, an indication of no embryonic lethality due to insertion location. All three lines were backcrossed to C57BL/6 ten times to establish congenic lines before analysis to eliminate genetic background that may contribute to variable expression patterns between lines and within the litters of each line. Genomic mapping analysis was performed on these lines using Targeted Locus Amplification (TLA) to identify the transgene integration sites and provide estimates on transgene copy number. The mouse mm10 genome was used as the reference sequence for alignment between transgene and host genome (Fig. 1B). The integration site for line pJ2-Cre:EGFP (#6) is at chromosome 11: 28042924 - 28617406. According to the reference sequence, the integration event affects the *Ccdc85a* (coiled-coil domain containing 85A) and *2810471M01Rik* gene regions. A complex integration has occurred here with different genomic rearrangements including transgene inversion, fusion, and deletion. The copy number is estimated to be 3-6 copies. The pJ2-Cre:EGFP (#7) 5’ integration site was determined to be chromosome 8: 41669071, which is near no annotated genes; however, the 3’ integration site could not be identified, suggesting it is within a repetitive or low complexity region, which are less efficiently sequenced and show low to no sequence coverage. The copy number is estimated to be 24-33 copies. The pJ2-Cre:EGFP (#11) transgene was integrated into chromosome 11: 42111411 - 42135537. The 24-kb genomic sequence between the two identified breakpoints is duplicated and present at both ends of the integrated sequence. The integration event and genomic duplication includes exons 9 and 10 of the *Gabra1* (Gamma-aminobutyric acid [GABA] A receptor, subunit alpha1) gene. The copy number is estimated to be 28-78 copies. To confirm these mapping analyses, we performed genomic PCR with primers between the transgene and host genome in each of the three lines and detected a PCR product of the expected size in all cases (Fig. 1C).

### *DUX4*-expressing cell lineages are present in the forelimb, hindlimb, and face during development

To visualize cell lineages where the *DUX4* enhancers and promoter have been active, we used *R26^NZG^* reporter mice (Yamamoto et al., 2009), which express a nuclear localized LacZ under the control of the ubiquitous CAG promoter in response to cre recombinase expression (Fig. S1A). Thus, in double transgenic mice, any cell in which the *DUX4* regulatory elements were active would express cre, leading to stable recombination of the reporter and expression of LacZ in the subsequent cell lineages, which is detected by X-gal staining at high sensitivity. Since some enhancers are active during oogenesis and Cre protein persists post meiotically in the oocyte, we first evaluated whether pJ2-Cre:EGFP has maternally inherited activity. We found that all embryos produced from female *pJ2-Cre:EGFP/+* mice crossed with male *R26^NZG/NZG^* reporter mice showed a recombined transgene in the amniotic sac and umbilical cord, and expressed ubiquitous LacZ regardless of inheritance of the pJ2-Cre:EGFP transgene (Fig. S1B-C). This indicates that the DUX4 regulatory elements are active in the female germline. Fortunately, embryos produced from female *R26^NZG^* crossed with male *pJ2-Cre:EGFP/+* mice did not show Cre activity in the umbilical cord or amniotic sac (Fig. S1D), indicating no paternal inheritance of Cre activity. Thus, all future litters in this study were generated using male *pJ2-Cre:EGFP* mice crossed with female *R26^NZG^* mice.

During development, myogenesis occurs in two stages: primary myogenesis in which Pax3+ progenitors arise from the dermomyotome to form multinucleated primary myofibers (embryonic stage E10-E12), and secondary myogenesis in which Pax7+ progenitors form secondary fibers by using primary fibers as a scaffold, thus contributing to the growth of fetal muscle (E14.5-E17.5) (Chal and Pourquie, 2017). To determine if the *DUX4* regulatory elements are active during embryonic myogenesis, *pJ2-Cre:EGFP/+; R26^NZG/+^* double transgenic embryos were generated from three independent-insertion lines, #6, #7 and #11, and reporter expression was analyzed at various stages. Intriguingly, X-gal-positive cells were commonly detected close to the dermis in both limbs and at the corner of the mouth in all three lines during E12.5 to E14.5, although the pattern, intensity, and timing of LacZ expression were variable (Figs. 2, S2 and S3). Since FSHD is a skeletal muscle disease and the *DUX4* transgene contains two myogenic enhancers, it was expected that expression would be found in skeletal muscle lineages. To determine the X-gal staining patterns for skeletal muscle lineages, *ACTA1-cre/+; R26^NZG/+^* embryos were analyzed (Figs. 2, S2 and S3). Surprisingly, all lines showed minimally overlapping staining with that of *ACTA1-cre/+; R26^NZG/+^* embryos, with the exception of line #6, which showed staining in facial expression muscles located close to the surface (Lescroart et al., 2010)(Figs. 2 and S3C-E). We identified the following X-gal-positive facial expression muscles: auricularis (au; connects the ear to the skull), buccinator (bu; used for chewing, sucking, and blowing), frontalis (fr; raises the eyebrow), orbitalis oculi (oo; closes the eyelid), quadratus labii (qua; moves the upper lip, used for smiling), and zygomaticus (zy; used for smiling). The facial expression muscles originate from mesodermal progenitor cells in the second branchial arch (BA2). We confirmed that line #6 *pJ2-Cre:EGFP/+; R26^NZG/+^* embryos showed X-gal-positive cells in BA2 at E10.5 (Fig. S3A), indicating that *DUX4* regulatory elements in line #6 are active in the branchiomeric muscle lineage that gives rise to facial expression muscles. Line #11 also showed similar but less consistent X-gal staining in the face (Figs. 2E and S2D), and with more ubiquitous staining in head and ventral trunk close to the surface (Fig. 2E, J, and O). It is possible that reporter expression in line #6 is influenced by gene regulation for the neighboring *Efemp1* gene, which is expressed in the 1^st^, 2^nd^ and 3^rd^ BA at the same stage (Ehlermann et al., 2003) thus resulting in a more extensive pattern of facial expression than the other two insertion lines.

**Figure 2.**
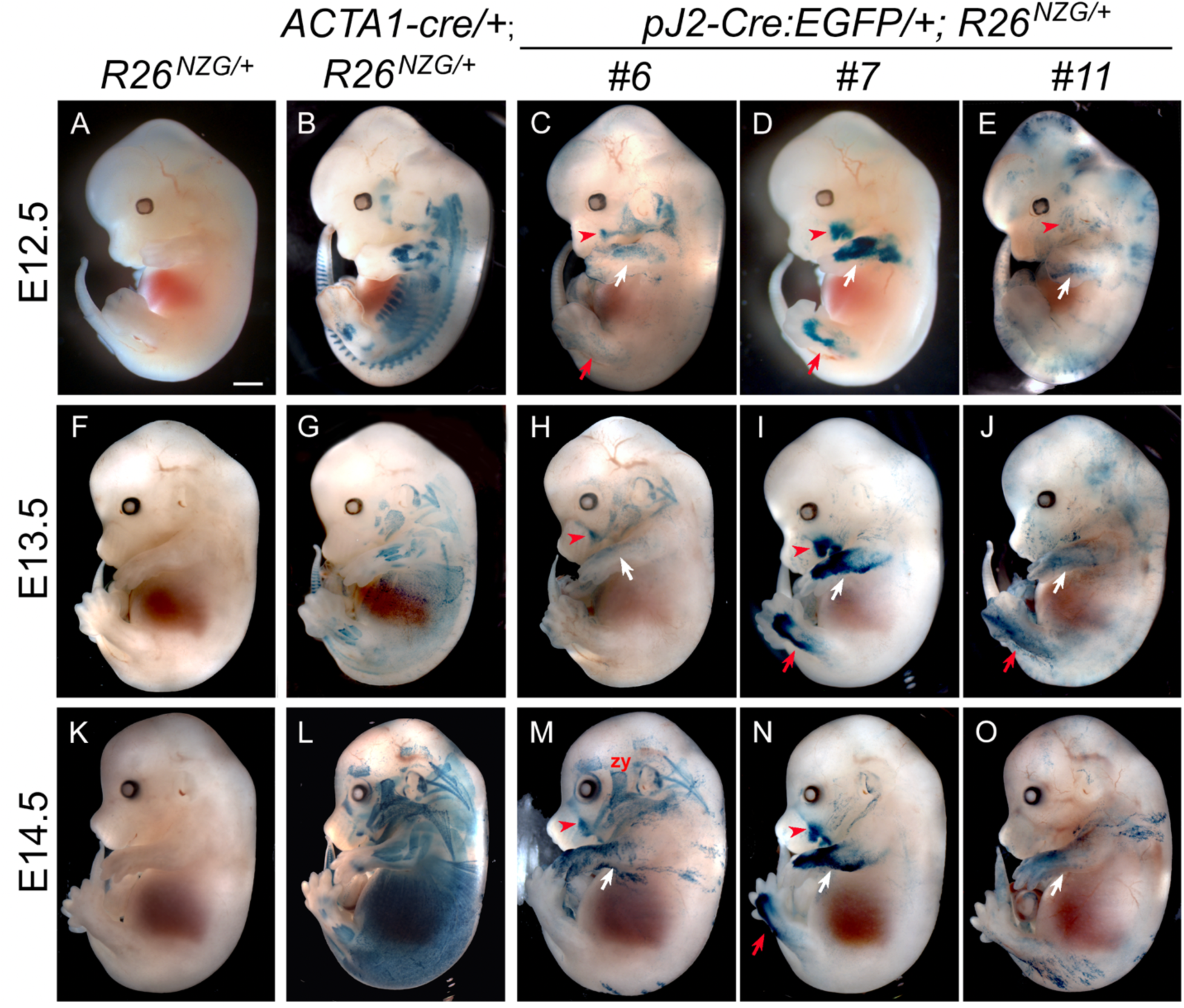
Activity of *DUX4* regulatory elements in limb and face during murine embryonic development. X-gal staining of *pJ2-Cre:EGFP/+; R26^NZG/+^* double transgenic embryos at E12.5 (C-E), E13.5 (H-J) and E14.5 (M-O). In all three independent insertion lines of transgene, #6, #7 and #11, DUX4 regulatory element is active in forelimb (white arrows), hindlimb (red arrows) and face where zygomaticus muscle (zy) connected at the corner of lip (red arrowhead). X-gal staining pattern of embryonic skeletal muscles in *ACTA1-cre/+; R26^NZG/+^* double transgenic embryos are shown at each embryonic stage as example of pan skeletal muscle staining (B, G, L) along with embryos with only reporter transgene (A, F, K) as negative control. Scale bar: 1mm.

Within each line, X-gal staining among littermates was more variable compared to *ACTA1-cre/+; R26^NZG/+^* embryos, with the exception of line #7 *pJ2-Cre:EGFP/+; R26^NZG/+^* which displayed the most consistent staining pattern, mainly in the limb and at the corner of the mouth (Fig. S2). Importantly, line #7, the only line in which the transgene is likely free of integration effects, displayed more consistent staining in the forelimbs, hindlimbs, and face throughout embryogenesis and within litters; this pattern is likely the truest indicator of cells derived from DUX4-positive lineages (Figs. 2D, I and N, S2C). To further evaluate the identity and location of X-gal positive cell lineages, we analyzed tissue sections from the forelimb of line #7 *pJ2-Cre:EGFP/+; R26^NZG/+^* E13.5 embryos by X-gal staining and Myosin heavy chain 1 (MYH1) immunostaining (Fig. 3). In the lower forelimb, X-gal-positive cells were observed in the dorsal mesenchyme adjacent to extensor muscles positive for MYH1 (Fig. 3C, D) just below the ectoderm (Fig. 3E, F). In the upper forelimb, X-gal staining was observed in the ventral mesenchyme next to the humerus and in the cells surrounding vascular structures (Fig. 3G, H). Overall, in the three insertion lines, cell lineages in which the *DUX4* regulatory elements are active are observed in the face and in the developing mesenchyme of the dorsal-anterior lower forelimb and lower hindlimb, but there is minimal overlap with embryonic myofibers.

**Figure 3.**
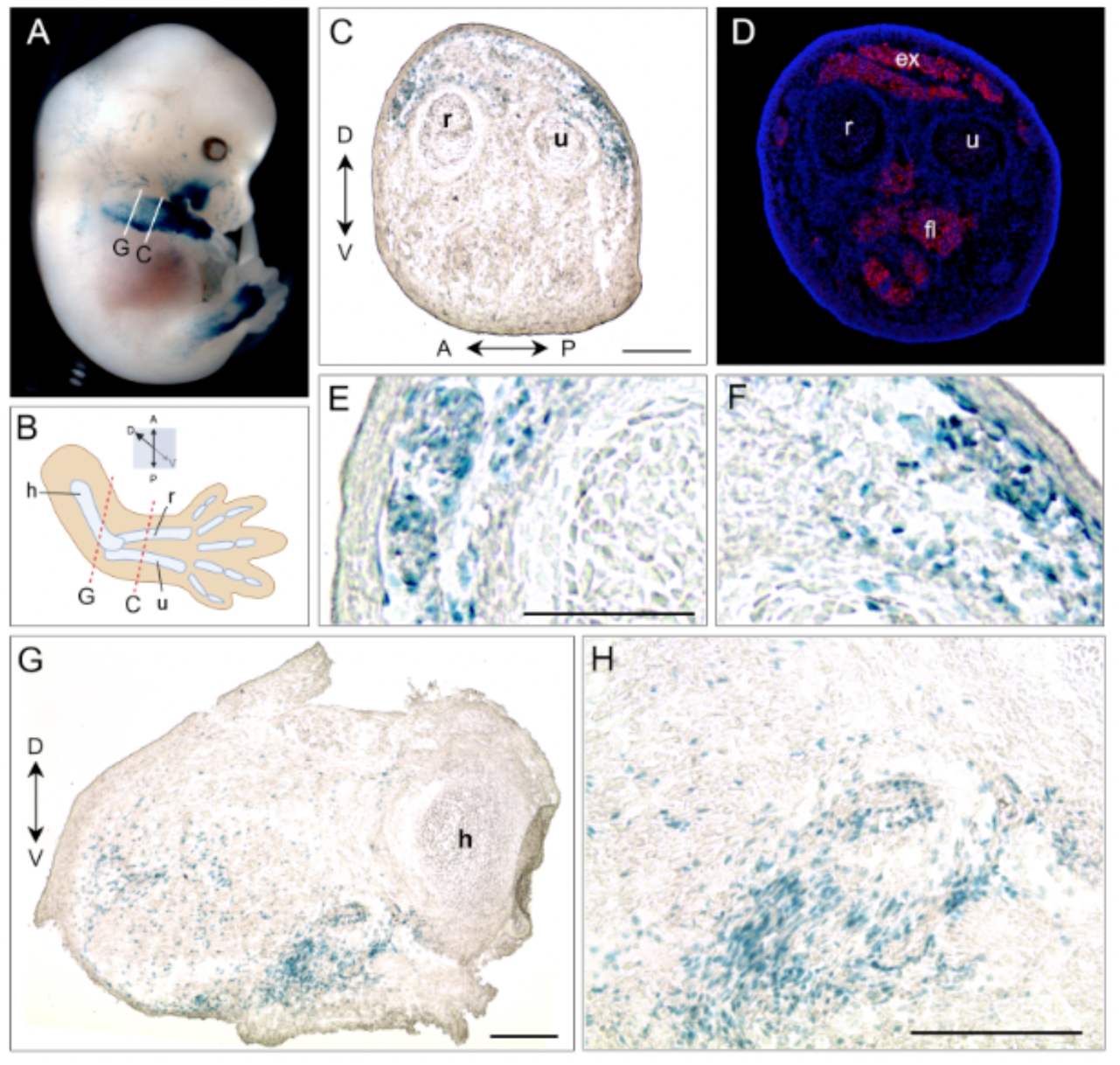
pJ2-Cre:EGFP #7/+; *R26^NZG/+^* embryos show very localized X-gal-positive cells in forelimb, hindlimb and side of mouth. (A) Right forelimb of X-gal stained E13.5 embryo was sectioned at place indicated with white lines. (B) Simplified illustration of forelimb with the radius (r), ulna (u) and humerus (h) bones, and anterior (A), posterior (P), dorsal (D) and ventral (V) directions. Sectioned planes are indicated in red dash lines. (C, E, and F) X-gal staining in subepidermal mesenchymal cells at dorsal of lower forelimb. (D) The serial section of C was stained with antibodies to myosin heavy chain 1 (MF20, red) and DAPI (blue). The extensor (ex) and flexor (fl) muscles are indicated. (G, H) X-gal signal in ventral mesenchyme of upper forelimb, and in cells surrounding vascular structures (arrows). Scale bar: 200µm.

### Adult tissues contain *DUX4*-expressing cell lineages

We next investigated the activity of *DUX4* regulatory elements in adult tissues using the same *R26^NGZ^* reporter mice. Since most X-gal-positive lineages in *pJ2-Cre:EGFP/+; R26^NGZ/+^*embryos were in the mesenchyme outside of embryonic myofibers, we analyzed non-muscle tissues (brain, thymus, heart, lung, liver, kidney, spleen, testis, and uterus) in addition to skeletal muscles (cheek, triceps, and tibialis anterior [TA]) of adult mice (Fig 4). Among these, testis was strongly X-gal-positive in all three independent-insertion lines (Fig. 4), confirming that the transgene constructs recapitulated what little is known about *DUX4* expression in adult tissues. X-gal staining of a testis cross-section showed that elongating spermatids at the late stage of spermatogenesis, close to the lumen in seminiferous tubules, were X-gal-positive (Fig. 5A-D). In addition, mature sperm isolated from *pJ2-Cre:EGFP/+; R26^NGZ/+^* males at ∼8 weeks of age were X-gal-positive in all three lines (Fig. 5E-H). For lines #6 and #11, the X-gal signals in skeletal muscle as well as thymus, heart, lung, spleen, and uterus were relatively weak (Fig. 4). Line-specific X-gal staining was observed in liver and kidney in line #6 *pJ2-Cre:EGFP; R26^NZG/+^* mice and part of the brain in line #11 *pJ2-Cre:EGFP; R26^NZG/+^* mice. The latter is potentially a result of integration near the *Gabra1* locus, since Gabra1 functions as a receptor for the GABA neurotransmitter in the central nervous system (Gilsoul et al., 2019).

**Figure 4.**
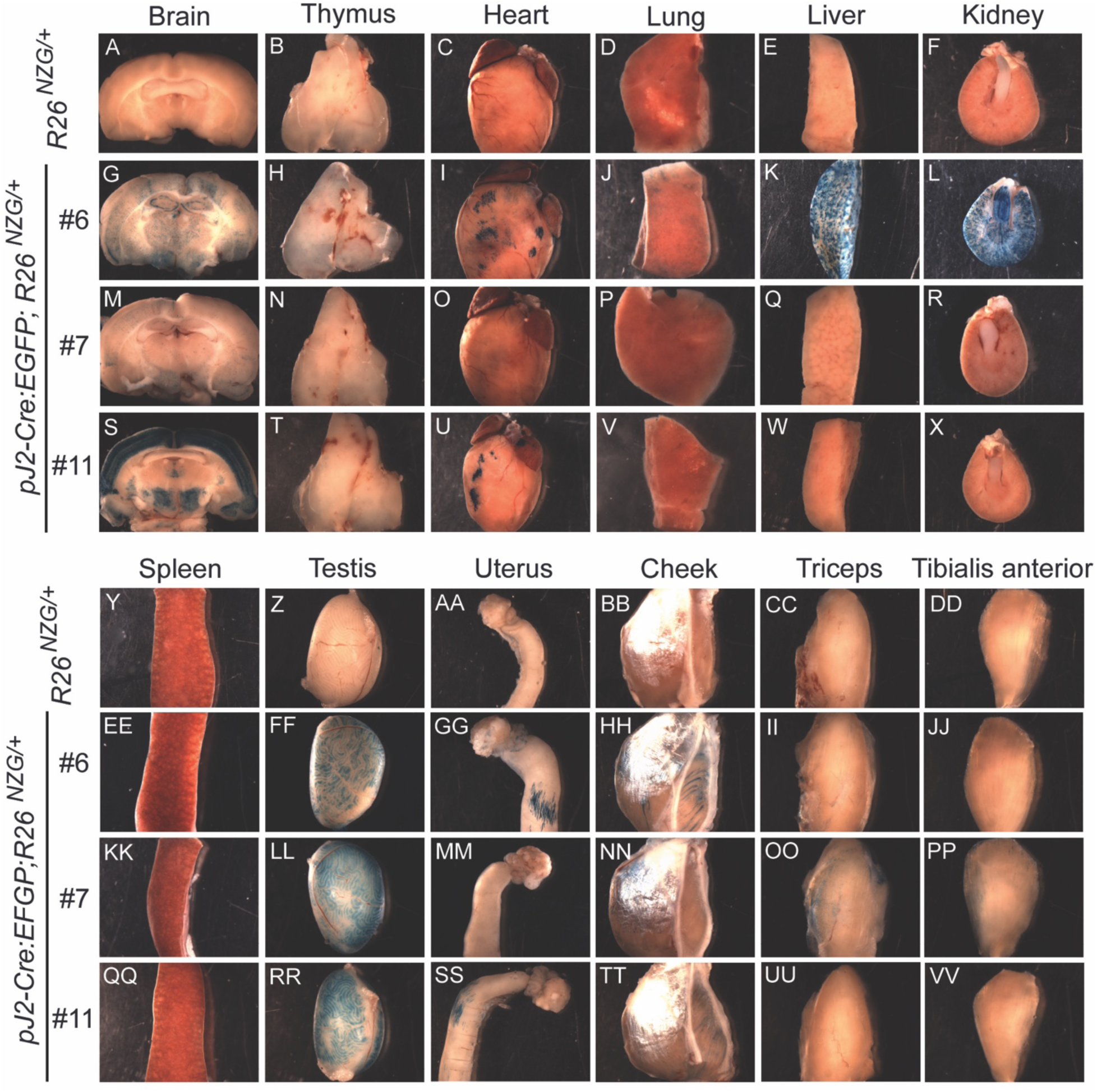
Activity of *DUX4* regulatory elements in adult murine tissues. X-gal staining for LacZ activity in *R26^NZG/+^*(A-F, Y-DD) and *pJ2-Cre:EGFP/+; R26^NZG/+^* lines #6 (G-L, EE-JJ), #7 (M-R, KK-PP), and #11 (S-X, QQ-VV) at >8 weeks of age in brain, thymus, heart, lung, liver, kidney, spleen, testis, uterus, and skeletal muscles (cheek, triceps, and tibialis anterior).

**Figure 5.**
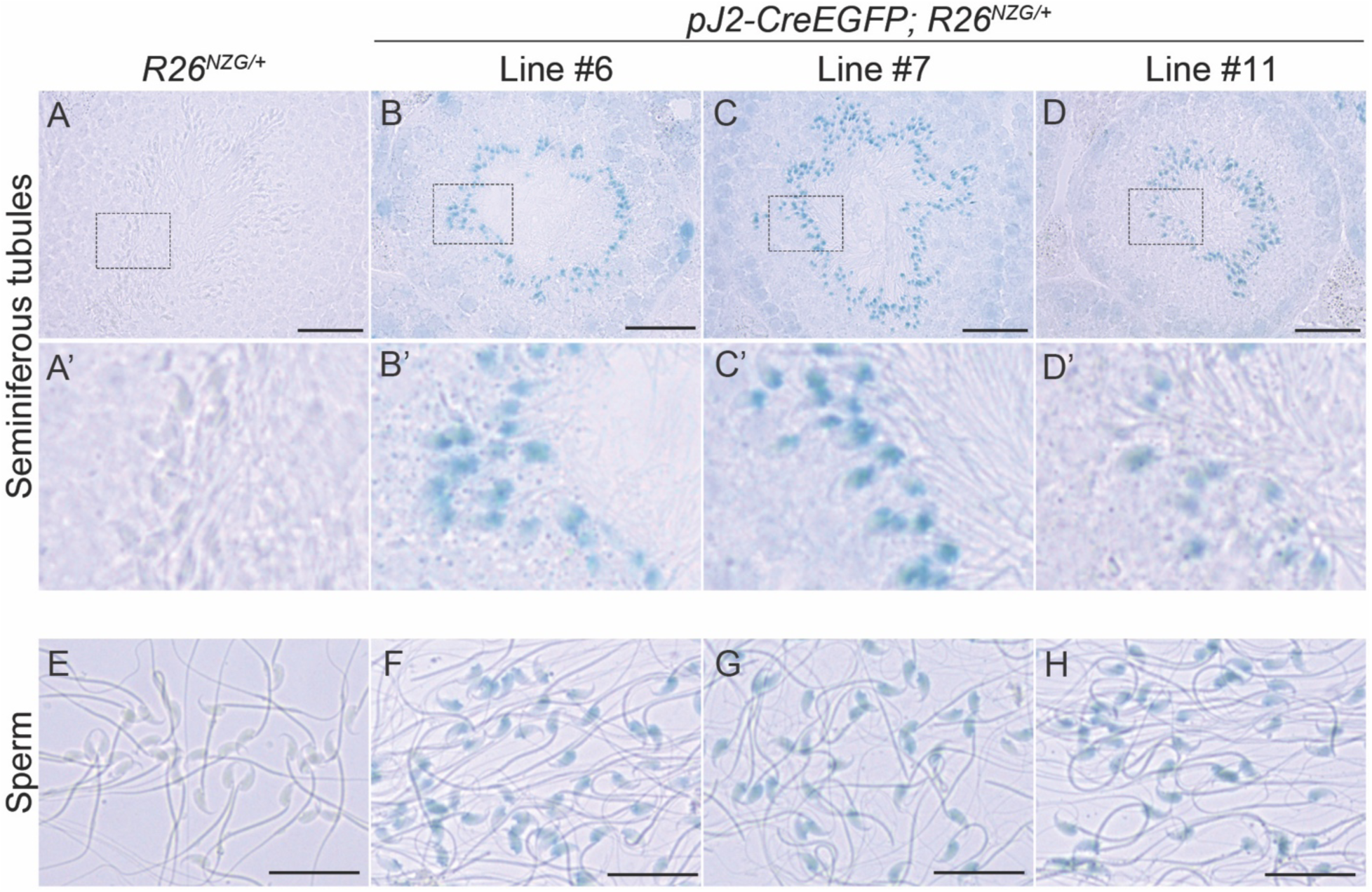
Activity of *DUX4* regulatory elements in male germ cells. X-gal staining for LacZ activity for *R26^NZG/+^* (A, A’, E) and *pJ2-Cre:EGFP/+; R26^NZG/+^* lines #6 (B, B’, F), line #7 (C, C’, G), and line #11 (D, D’, H) at >8 weeks of age in a cross-section of testis (A-D). The middle panel (A’-D’) shows the enlargement of a rectangle found in the upper panel of the respective figure. Scale bar = 50 μm. Lower panel (E-H) shows matured sperm from ∼8 weeks old mice. Scale bar = 25 μm.

Line #7, which displayed consistent X-gal staining in the embryonic limbs and face, showed weak X-gal staining in adult muscles and strong X-gal staining in testis, without apparent staining in other non-muscle tissues, recapitulating what is known about *DUX4* expression in FSHD patients.

### Pericytes are a *DUX4*-expressing cell lineage

Although the *DUX4* DMEs are known to be active in human and murine myogenic cells (Himeda et al., 2014), we did not see a strong X-gal signal in whole skeletal muscle mounts, which is consistent with rare expression of DUX4 in FSHD skeletal muscle (Jones et al., 2012; Tassin et al., 2013). However, cross-sections of TA muscles showed the presence of X-gal-positive cells in all three *pJ2-Cre:EGFP; R26^NZG/+^* transgenic lines (Fig. 6). While X-gal-positive cells were detected within myofibers (Fig. S4), they were also located outside of myofibers and within the interstitial space (Fig. 6). Interestingly, in all three *pJ2-Cre:EGFP; R26^NZG/+^* transgenic lines, X-gal-positive cells were often located near blood vessels in TA muscles (Fig. 6). Since DUX4 expression is causal for FSHD (Lemmers et al., 2010; Snider et al., 2010), and FSHD pathology is associated with active muscle regeneration (Snider et al., 2010), we investigated whether the *DUX4* regulatory elements were active during skeletal muscle regeneration. Skeletal muscle injury was induced in *pJ2-Cre:EGFP; R26^NZG/+^*mice by injecting barium chloride (BaCl_2_) into the TA muscle and then allowing regeneration to occur. At 10 days post-injury, the centralized myonuclei of regenerating myofibers were found to be X-gal-positive (Fig. 6G-L and Fig S5).

**Figure 6.**
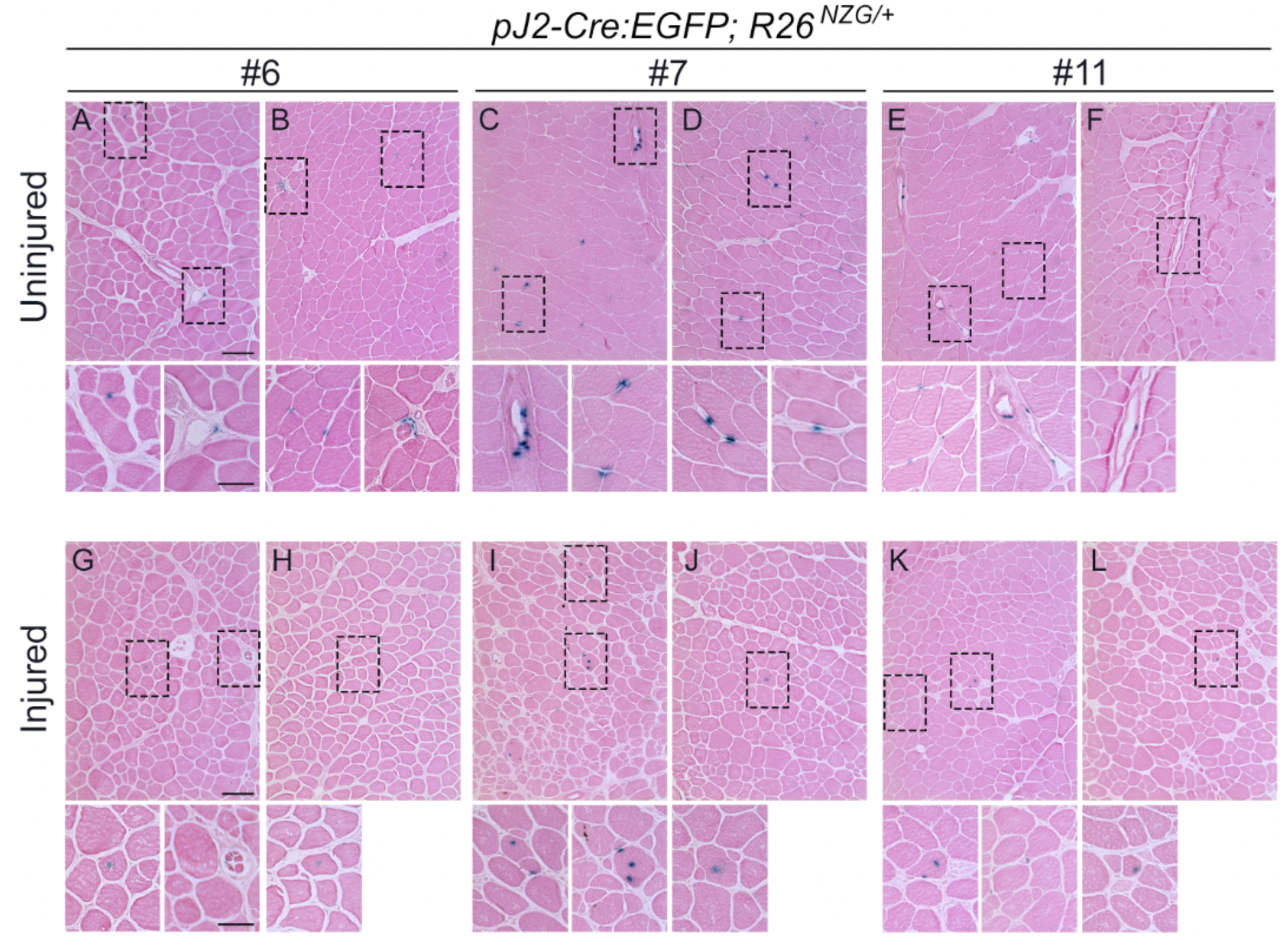
Activity of *DUX4* regulatory elements in adult skeletal muscle. (A-F) X-gal and eosin staining of TA muscle sections from two animals for each *pJ2-Cre:EGFP/+; R26^NZG/+^* lines; #6 (A, B), #7 (C, D), and #11 (E, F) at >8 weeks of age. Scale bar in A=100 μm. Interstitially localized X-gal-positive cells in A-F were magnified below. Sale bar =50 μm. (G-L) X-gal and eosin staining of injured TA muscles at 10 days after barium chloride injection. Two animals for each *pJ2-Cre:EGFP/+; R26^NZG/+^* lines; #6 (G, H), #7 (I, J), and #11 (K, L) at >8 weeks of age. All X-gal-positive cells had centralized myonuclei. No X-gal signals were observed in *R26^NZG/+^* TA muscles in Fig S5.

This data suggested that DUX4-positive cell lineages may be involved in muscle regeneration. Myogenic satellite cells are the major cell type contributing to muscle repair and regeneration (von Maltzahn et al., 2013; Wang and Rudnicki, 2011). The transcription factor Pax7 is expressed in quiescent and activated muscle satellite cells and required for their function in adult skeletal muscle (Seale et al., 2000; von Maltzahn et al., 2013; Zammit et al., 2002). In addition, PAX7 target gene repression correlates with FSHD disease status (Banerji, 2020). To determine if X-gal-positive lineages in skeletal muscle from adult *pJ2-Cre:EGFP; R26^NZG/+^* mice include satellite cells, muscle sections were co-immunostained for Pax7 and beta-galactosidase (Figs. 7 and S6). As expected, cells expressing either Pax7 or beta-galactosidase were rare. Regardless, no cells showed staining for both. All detected Pax7-positive cells were negative for beta-galactosidase, and more importantly, all detected beta-galactosidase-positive cells were negative for Pax7. Thus, while we cannot exclude the possibility that some fraction of Pax7-positive cells could be X-gal-positive as well (but undetected in our analysis), our overall data indicate that the *DUX4* regulatory elements are active, not in satellite cells, but in a lineage of interstitial cells.

**Figure 7.**
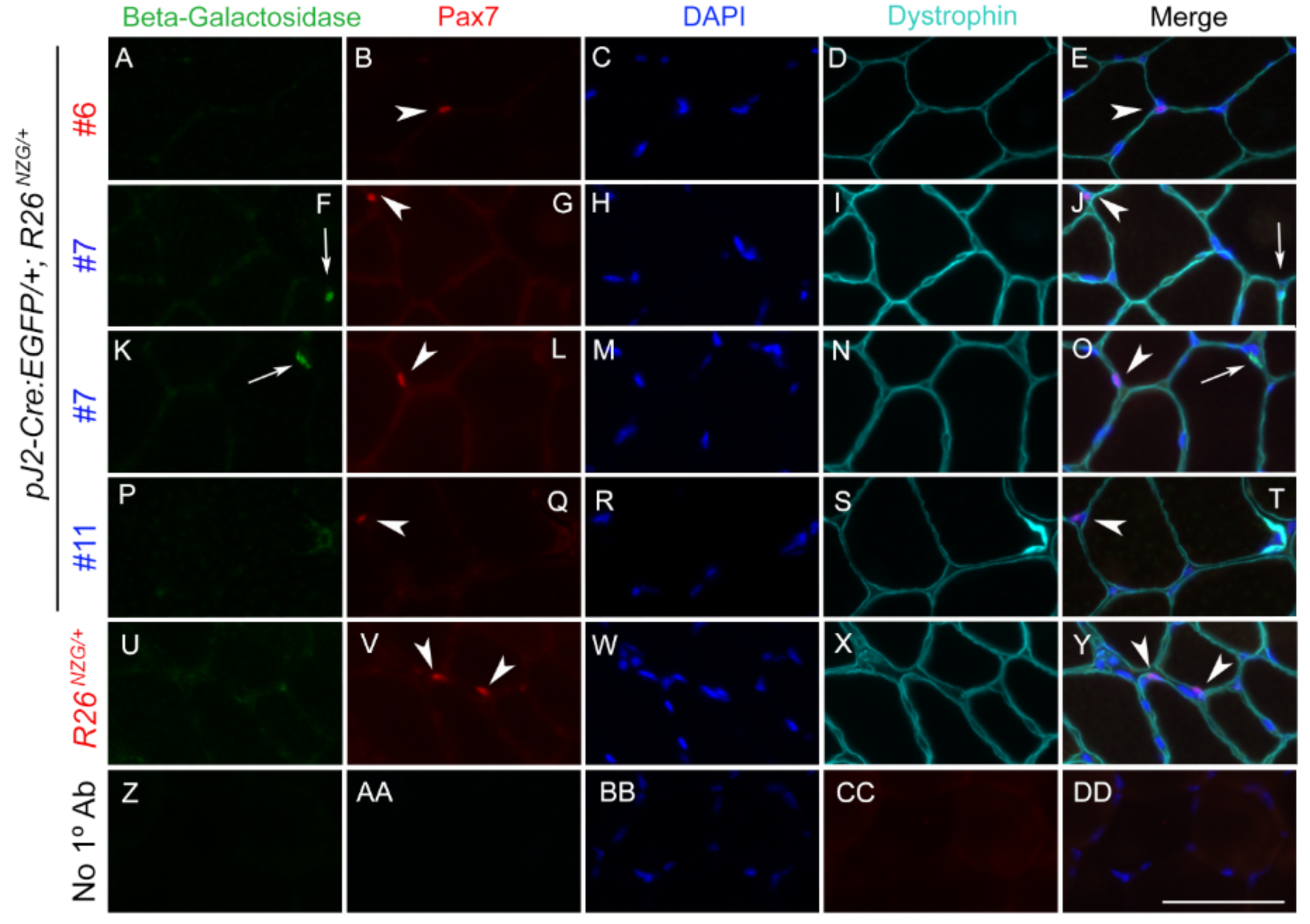
The interstitially localized cells in which *DUX4* regulatory elements are active are not Pax7+ muscle satellite cells. TA muscle sections from male (blue) and female (red) *pJ2-Cre:EGFP/+; R26^NZG/+^* and *R26^NZG/+^* mice were immunostained for beta-galactosidase (green, A, F, K, P, U), Pax7 (red, B, G, L, Q, V), and dystrophin (aqua, D, I, N, S, X) and stained with DAPI (blue, C, H, M, R, W, BB) to show nuclei. Z-DD) no primary antibody control. White arrows show beta-galactosidase positive nuclei, white arrowheads show Pax7 positive nuclei. Scale bar = 50 μm.

Pericytes, which are myogenic precursor cells distinct from myogenic satellite cells, are one of the cell types found in the small vessels of the skeletal muscle interstitial space (Dellavalle et al., 2007). To determine if the X-gal-positive interstitial cells in skeletal muscles of the *pJ2-Cre:EGFP/+; R26^NZG/+^* mice could be pericytes, skeletal muscle sections were stained for tissue nonspecific alkaline phosphatase (AP), a marker for pericytes and endothelial cells (Dellavalle et al., 2011). Indeed, some X-gal-positive cells in healthy muscles were positive for AP (Fig. 8A). Since AP is expressed in both pericytes and endothelial cells, we stained for the pericyte marker *Cspg4* (also known as NG2) and found that some X-gal-positive cells were surrounded by the CSPG4 signal (Fig. 8B). This data indicates that the *DUX4* regulatory regions were active at some point within the pericyte cell lineage.

**Figure 8.**
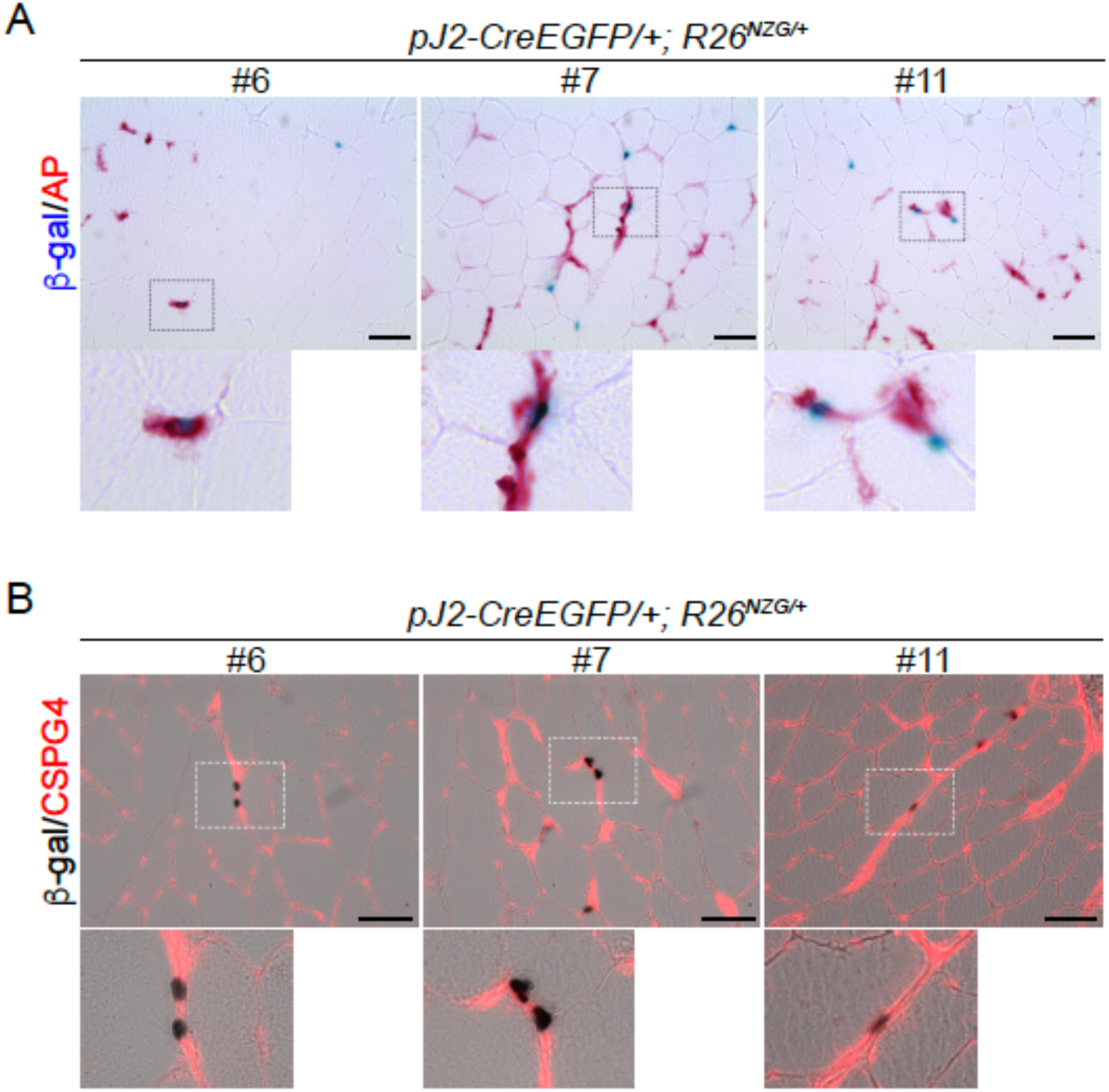
*DUX4* regulatory elements are active in the pericyte lineage. A) X-gal and alkaline phosphatase (AP) staining and B) X-gal and immunofluorescence staining for CSPG4 of normal TA muscle sections at >8 weeks of age. Scale bar: 50 μm. The lower panel shows the enlargement of the rectangle in the upper panel.

## Discussion

The goal of this study was to identify the tissues and cell lineages where the human *DUX4* enhancer/promoter regulatory elements are active in mice. The scientific rationale for generating *pJ2-Cre:EGFP* transgenic mice was that although the human *DUX4* gene is primate-specific (Leidenroth et al., 2012; Leidenroth and Hewitt, 2010), many regulatory elements, pathways, and transcription factors are conserved between mice and humans (Cheng et al., 2014; Diehl and Boyle, 2018). Interestingly, the *DUX* family genes show high conservation of expression and function with respect to zygotic genome activation across species (De Iaco et al., 2017; Hendrickson et al., 2017; Whiddon et al., 2017) and are encoded within D4Z4-like repeats, although the size of each RU and array varies (Clapp et al., 2007). Thus, the signals and trans-acting factors that regulate *DUX4* expression in humans with and without FSHD could be similarly conserved in mice. In fact, our initial work that identified and characterized the *DUX4* myogenic enhancers showed that they were similarly active in both human and murine myogenic cells (Himeda et al., 2014), further validating this approach. In most humans that do not have FSHD, *DUX4* is present in an extended D4Z4 repeat array and is epigenetically silenced and thus not expressed in somatic tissues, whereas in FSHD1, the D4Z4 array is contracted and *DUX4* is epigenetically de-repressed (de Greef et al., 2009; Himeda and Jones, 2019; Snider et al., 2010; van Deutekom et al., 1993; van Overveld et al., 2003; Wijmenga et al., 1992). Therefore, the *pJ2-Cre:EGFP* transgenic mice, which contain a single D4Z4 RU as found in severe FSHD1, when crossed with *R26^NZG^* reporter mice, allowed us to visualize the activity of the *DUX4* regulatory elements in vivo under FSHD-like epigenetic conditions, and should be a useful tool for studying factors and stimuli that affect regulation of *DUX4* expression in FSHD.

In vivo regulation of *DUX4* within the context of a D4Z4 repeat array has previously been studied to some degree in the first published FSHD mouse models, the D4Z4-2.5 and D4Z4-12.5 mice, which differentially contained some of the human *DUX4* regulatory elements (Krom et al., 2013). The D4Z4-2.5 mouse transgene contains a 13.5-kb EcoRI restriction fragment spanning a contracted chromosome 4q35 2.5RU D4Z4 array isolated from an FSHD patient and includes DME1 (but not DME2) and the FSHD-permissive 4A subtelomere, but lacks the downstream developmental noncoding exons 6 and 7 with the developmentally utilized PAS (Gabriels et al., 1999; Snider et al., 2010). The D4Z4-12.5 mouse has significant additional centromere proximal sequence containing the upstream *FRG1* and *FRG2* genes and includes DME1, DME2, and the FSHD-permissive 4A subtelomere. While this mouse lacks the downstream developmental exons 6 and 7 and PAS, the healthy-sized 12.5 D4Z4 RU array causes epigenetic repression of the *DUX4* transgene (Krom et al., 2013). In comparison, the *pJ2-Cre:EGFP* mice used here have four key differences in transgene design. First, the pJ2-CreEGFP mouse has a single D4Z4 RU, albeit only the 1817 bp upstream of the *DUX4* open reading frame, with both DME1 and DME2. Thus, the genetic regulation is comparable to the non-FSHD 12.5-D4Z4 mouse, with the exception that the pJ2-CreEGFP mouse is in the FSHD1 genetic state (1 D4Z4 RU) and thus not epigenetically repressed. Second, the Cre:EGFP fusion gene is in place of the *DUX4* gene and therefore cre expression will genetically mark cells and their lineages instead of possibly killing the cells (Kowaljow et al., 2007). Third, the Cre:EGFP reporter gene utilizes a β-globin PAS instead of the FSHD-permissive PAS sequence, while the 2.5-D4Z4 and 12.5-D4Z4 mice are lacking the developmental PAS, which may or may not have consequences on developmental expression profiles. Finally, the transgene integration sites are different for all the mouse models, including chromosome 17 for the D4Z4-2.5 transgene and chromosome 2 for D4Z4-12.5 transgene (Krom et al., 2013). It should be noted that only single lines of both the D4Z4-2.5 and −12.5 mice were developed and analyzed, despite being generated by random insertion. Therefore, the effects of the integration site on the D4Z4 transgene regulation are not known. Overall, the pJ2-Cre:EGFP mice provide a unique opportunity to investigate developmental *DUX4* expression in the FSHD1-like state. Of note, the initial proof-of-concept analysis identified both murine testis and skeletal muscle tissue (Figs 2-5) as positive for *DUX4* expression, confirming what is known about *DUX4* expression in adult humans with and without FSHD (Jones et al., 2012; Snider et al., 2010; Yao et al., 2014).

### Human *DUX4* enhancer/promoter activity in mice

Comparing reporter expression among adult tissues of *pJ2-Cre:EGFP; R26^NZG^/+* mice, the testis, especially its germ cells, was strongly X-gal-positive in all three transgenic mouse lines. These results are similar to the abundant expression of *DUX4* found in testis of both the D4Z4-2.5 and D4Z4-12.5 mice (Krom et al., 2013) and humans (Snider et al., 2010). Moreover, the *DUX4* expression in D4Z4-2.5 mice is observed in cells near the periphery of the seminiferous tubules, likely in spermatogonia and primary spermatocytes. The higher amount of *DUX4* mRNA in testis and the expression in spermatogonia or spermatocytes was also confirmed in human testis (Snider et al., 2010), indicating that the regulatory elements we used function similarly in murine testis as they do in the human tissue.

With respect to somatic tissues, the *pJ2-Cre:EGFP; R26^NZG^/+* adult mice showed some variable and line-specific tissue expression patterns outside of skeletal muscle (Fig. 4); however, double transgenic mice from all three lines exhibited positive, albeit very limited expression in skeletal muscles. X-gal-positive staining was observed in the interstitial space, within myofibers, and in the centralized myonuclei of regenerating myofibers. In FSHD, despite all of the cells sharing the same genetic defect and similar epigenetic dysregulation of the pathogenic D4Z4 array and *DUX4* gene, it is extremely rare to find DUX4-positive cells in muscle biopsies from patients, and even cultured FSHD myotubes show only ∼1/1000 DUX4-positive myonuclei (Jones et al., 2012; Rickard et al., 2015; Tassin et al., 2013). In fact, the only real evidence that DUX4 protein is present in FSHD patient muscles is the misexpression of DUX4 target genes, which serves as a surrogate of DUX4 activity (Tawil et al., 2024; Wong et al., 2024; Wong et al., 2020; Yao et al., 2014). Thus, the skeletal muscles from the adult *pJ2-Cre:EGFP; R26^NZG^/+* mice exhibiting rare X-gal-positive nuclei, despite all the cells in the tissue containing the same D4Z4 transgene and reporter gene, mimic the rare DUX4 protein expression found in FSHD muscles. While it is assumed that DUX4 expression within FSHD muscles occurs in myofibers, this will likely require single cell RNA sequencing analysis to correlate DUX4 gene expression signatures with cell-specific signatures. The work presented here (Figs 7, 8, and S6) suggests that PAX7-positive satellite cells are unlikely candidates for expressing DUX4, while pericytes may be a novel DUX4-expressing cell lineage in FSHD muscle.

Considering that testis showed by far the most high and uniform X-gal staining that was consistent among mice (Figs. 4 and 5), it is likely that the DMEs contain elements that drive expression in testis, in addition to myogenic factor binding motifs. DME2 also likely contains elements that prevent expression in some non-myogenic cell types, such as fibroblasts (Himeda et al., 2014). In comparison, the somatic tissues of the D4Z4-2.5 mice (which contain DME1, but not DME2) showed fairly reproducible levels of *DUX4* mRNA throughout the body, including in skeletal muscles (limbs, trunk, and head) and all non-muscle tissues tested (except for the liver, where expression was more variable). Thus, the pJ2-Cre:EGFP mouse serves as a better model than the D4Z4-2.5 mouse for developmental studies of DUX4 regulation and expression. In contrast, due to epigenetic repression of the extended array, the D4Z4-12.5 mice showed rare and sporadic *DUX4* expression in only a few skeletal muscles (TA and pectoralis muscles), while all other somatic tissues were generally silent or showed inconsistent *DUX4* expression (Krom et al., 2013). Since the genetics and epigenetics of the model resemble those of the non-FSHD state, this would also not be a good model for developmental studies investigating the role of DUX4 in FSHD.

This cell lineage study examining the activity of *DUX4* regulatory elements during development sheds light on the poorly understood spatiotemporal pattern of DUX4 expression in FSHD. Three independent-insertion lines of the transgene pJ2-Cre:EGFP produced X-gal staining in dorsal-anterior mesenchyme of limbs and a localized signal at the corner of the mouth, overlapping the general areas of clinical presentation in FSHD patient muscles. Transgene integration effects are a concern for two of our lines (#6 and #11); however, line #7, the only line in which the transgene is likely free of integration effects, displayed relatively consistent X-gal staining in the developing forelimbs, hindlimbs, and face. Since we didn’t observe GFP expression during ∼E10-E14.5 (data not shown), the *DUX4* regulatory elements are likely active early in embryogenesis and not constitutively active over the course of development. Interestingly, DUX4 has been shown to activate expression of H3.X and H3.Y, which are then incorporated into the bodies of other DUX4 target genes, priming them for enhanced re-activation in response to a second burst of DUX4 expression (Resnick et al., 2019). Thus, early embryonic expression of DUX4 may establish epigenetic marks that contribute to the postnatal activation of target genes in the limbs and face, leading to disease progression. Unfortunately, the reporter lines in the present study cannot be used to investigate this theory, as the H3.X and H3.Y variants are primate specific.

### Pericytes have a DUX4-positive lineage

Skeletal muscle histology from all three lines of *pJ2-Cre:EGFP; R26^NZG^/+* adult mice showed X-gal-positive cells located both within myofibers and in the interstitial space near blood vessels. We detected the presence of X-gal-positive centralized myonuclei in regenerating fibers following muscle injury; however, immunostaining for Pax7 and beta-galactosidase failed to identify any X-gal-positive satellite cells (Figs 7 and S6). While we cannot rule out the possibility that a small population of X-gal-positive satellite cells exists that were missed in our limited analysis, our results suggest that DUX4 may be expressed in non-muscle cells residing within the skeletal muscle that contribute to regeneration. Indeed, X-gal-positive interstitial cells were found to express pericyte cellular markers (Fig 8). Pericytes, which arise from a distinctly different lineage than muscle satellite cells, also contribute to skeletal muscle regeneration (Dellavalle et al., 2011; Dellavalle et al., 2007). Taken together, our findings suggest that DUX4 regulatory elements are active in the pericyte lineage and that this lineage may have a previously unknown role in FSHD. Considering that DUX4 expression is detrimental to muscle development and often toxic to somatic cells (Bosnakovski et al., 2017; Bosnakovski et al., 2018; Kowaljow et al., 2007; Mitsuhashi et al., 2013; Rickard et al., 2015; Tassin et al., 2013; Wallace et al., 2011; Wuebbles et al., 2010; Yao et al., 2014), this suggests the possibility that, in FSHD, aberrant expression of DUX4 in the pericyte developmental lineage might adversely impact the pericyte cell population and/or function, potentially contributing to FSHD pathophysiology over time.

### New tools for FSHD research

Overall, these novel transgenic mouse models with human *DUX4* regulatory elements are potentially a powerful new tool for investigating the underlying causes of FSHD pathology. For example, this initial work has: 1) identified a blood vessel-associated cell lineage that had, at some point in its developmental history, activated DUX4 expression, and 2) implicated the pericyte lineage as a novel source of developmental DUX4 expression that could impact skeletal muscle formation, growth, repair, and regeneration, thus potentially playing a role in FSHD pathology. In addition, the rare presence of X-gal-positive myonuclei in skeletal muscle recapitulates the rare mosaic DUX4 expression in FSHD skeletal muscle. As discussed earlier, the D4Z4 repeat in pJ2-Cre:EGFP mice mimics the situation in FSHD, where the 4q35 D4Z4 array is epigenetically dysregulated in all cells and cell types (de Greef et al., 2009; Jones et al., 2014); however, despite the loss of this repression, only a small fraction of FSHD skeletal muscle cells express DUX4 at any given time (Haynes et al., 2018; Jones et al., 2012). Interestingly, the fraction of DUX4-positive FSHD cells in culture can be significantly altered when the cells are subjected to certain stresses (Jones et al., 2015; Teveroni et al., 2017). One potential explanation is that DUX4 may be a stress and/or hormone responsive gene and FSHD pathology is induced by currently unknown signals that cause short-term bursts of DUX4 expression in increasing cell numbers, leading to the rapid accumulation of muscle pathology (Lim et al., 2020; Rickard et al., 2015). As such, the *pJ2-Cre:EGFP; R26^NZG^* mouse model could be used to identify factors, conditions, or stimuli that induce *DUX4* expression and, conversely, factors that prevent *DUX4* expression in vivo. Thus, our developmental models open up new in vivo discovery and therapeutic validation opportunities for understanding and treating FSHD.

## Material and methods

### Animals

All animal procedures were approved by the University of Nevada, Reno IACUC (Protocol #0701). Euthanasia was performed using CO_2_ followed by cervical dislocation. pJ2-Cre:EGFP mice were generated by the Jones lab at the University of Massachusetts Medical School transgenic mouse facility. ACTA1-cre mice (strain #006149) (Miniou et al., 1999) and *R26^NZG^* mice (stock #012429) (Yamamoto et al., 2009) were purchased from The Jackson Laboratory. Genotyping primers for cre recombinase are listed in Table S1. For the skeletal muscle injury procedure, 60 - 100 μl 1.2% barium chloride was injected into the right TA muscle with 31G needle under isoflurane anesthesia.

### Transgene construction

The pJ2-Cre:EGFP transgene was generated by digesting the pJ2 plasmid (Himeda et al., 2014) with FseI and AscI to remove the *DUX4* coding sequence. The replacement sequence was synthesized from FseI to the *DUX4* MAL start codon followed by the ATG and coding sequence for Cre:EGFP and the ß-globin polyadenylation signal (PAS) termination cassette, based on the published pCAG-Cre:GFP sequence (Matsuda and Cepko, 2007), with an AscI restriction site added at the 3’ end. This FseI/AscI fragment was ligated to the similarly digested pJ2 vector to create pJ2-Cre:EGFP and validated by sequencing.

### Mapping transgene integration sites

For identification of transgene integration sites in pJ2-Cre:EGFP mice, viable frozen mouse spleen cells (pJ2-Cre:EGFP #6) and bone marrow (pJ2-Cre:EGFP #7 and #11) were used and processed by Cergentis using the targeted locus amplification (TLA) protocol (de Vree et al., 2014). Two primer sets to the transgene were designed and used in individual TLA amplifications (Table S1). PCR products were purified, library prepped using the Illumina Nextera flex protocol and sequenced on an Illumina sequencer. Reads were mapped using BWA-SW, version 0.7.15-r1140, settings bwasw-b7 (Li and Durbin, 2010). The sequencing reads were aligned to the transgene sequence and the mouse mm10 genome was used as the host reference genome sequence. The PCR primers for confirming the integration sites are listed in Table S1.

### X-gal staining

A Leica CM1950 cryostat was used for making 10 μm cross-sections from skeletal muscle and testis frozen in liquid nitrogen-cooled isopentane. Fixation was performed in a solution of 2% paraformaldehyde (PFA), 0.25% glutaraldehyde, and 0.05% NP 40 for 15 minutes when using a cryosection,1-2 hours for embryos, and 1 hour for adult tissues, followed by washing 3 times in 1X PBS. Samples were then immersed in X-gal solution (1 mg/ml X-gal, 5 mM potassium ferricyanide, 5 mM potassium ferrocyanide, and 2 mM MgCl_2_) at 37°C for 45 – 120 minutes. Sperm smears derived from cauda epididymis were immersed in X-gal solution 37°C for 30 minutes without fixation. For X-gal plus eosin staining, after X-gal staining for 1 hour, the cross-sections were immersed in eosin solution for 10 seconds followed by 70-100 % ethanol and xylene. For X-gal plus alkaline phosphatase (AP) staining, cross-sections were fixed with 4% PFA for 10 minutes and immersed in X-gal solution at 37°C for 1 hour and then followed by PermaRed/AP (Diagnostic Biosystems, K049) for 10 minutes at room temperature.

For X-gal plus CSPG4 immunostaining, the cross-sections were first immersed in X-gal solution at 37°C for 1 hour and then followed by immunostaining. Fixation was performed in 4% PFA for 10 minutes followed by treatment with 0.25% TritonX-100 for 15 minutes and blocking solution (5% normal goat serum and 0.01% TritonX-100) for 30 minutes. The primary anti-CSPG4 antibody (Millipore, AB5320: 1:200) was incubated at 4°C overnight and then secondary antibody (Alexa 594 goat anti rabbit IgG, Invitrogen, A11037: 1:400) at room temperature for 1 hour.

### Immunofluorescence

The cross-sections were fixed in 4% PFA for 10 minutes followed by treatment with 100 mM glycine for 10 min, 0.25% TritonX-100 for 20 minutes and blocking with M.O.M. Immunodetection kit (Vector Laboratories) for 30 minutes. The primary antibodies (Beta Galactosidase (ICL, #CGAL-45A, 1:200), Pax7 (DSHB, 1:5), dystrophin (abcam, ab15277, 1:200) and MYH1 (DSHB, MF20, 1:5)) were incubated at 4°C overnight and then secondary antibodies (Goat anti-chicken IgY DyLight 488 (ThermoFisher, SA5-10070, 1:300), Alexa 594 donkey anti-mouse IgG (Jackson ImmunoResearch, 715-586-151, 1:300), and Alexa 647 donkey anti-rabbit (Jackson ImmunoResearch, 711-606-152, 1:300)) were incubated at room temperature for 1 hour. ProLong Gold Antifade Mountant with DAPI (ThermoFisher) was used for staining nuclei.

### Imaging

For images of sperm and a cross-section of skeletal muscle and testis, a Leica DM 2000 LED microscope, DFC290 camera, and LAS V4.12 software were used. For images of embryo and adult tissue, a ZEISS Stemi 2000-C microscope, Qimaging MP3.3-RTV-CLR-10 camera, and Qcapture suite software were used. For images of embryo at E14.5, two images were merged by photoshop. For images including immunofluorescence, a LEICA DMi8 microscope, DFC365 FX camera, and LAS X software were used.

## Author contributions statement

YH, PLJ, and TIJ conceived of the study. YH and TIJ performed experiments and collected data. YH, CLH, PLJ, and TIJ analyzed data and wrote the manuscript.

