## Supplementary material for "Transgenic mouse models for investigating human *DUX4* expression during development and its roles in FSHD pathophysiology": Fig. S

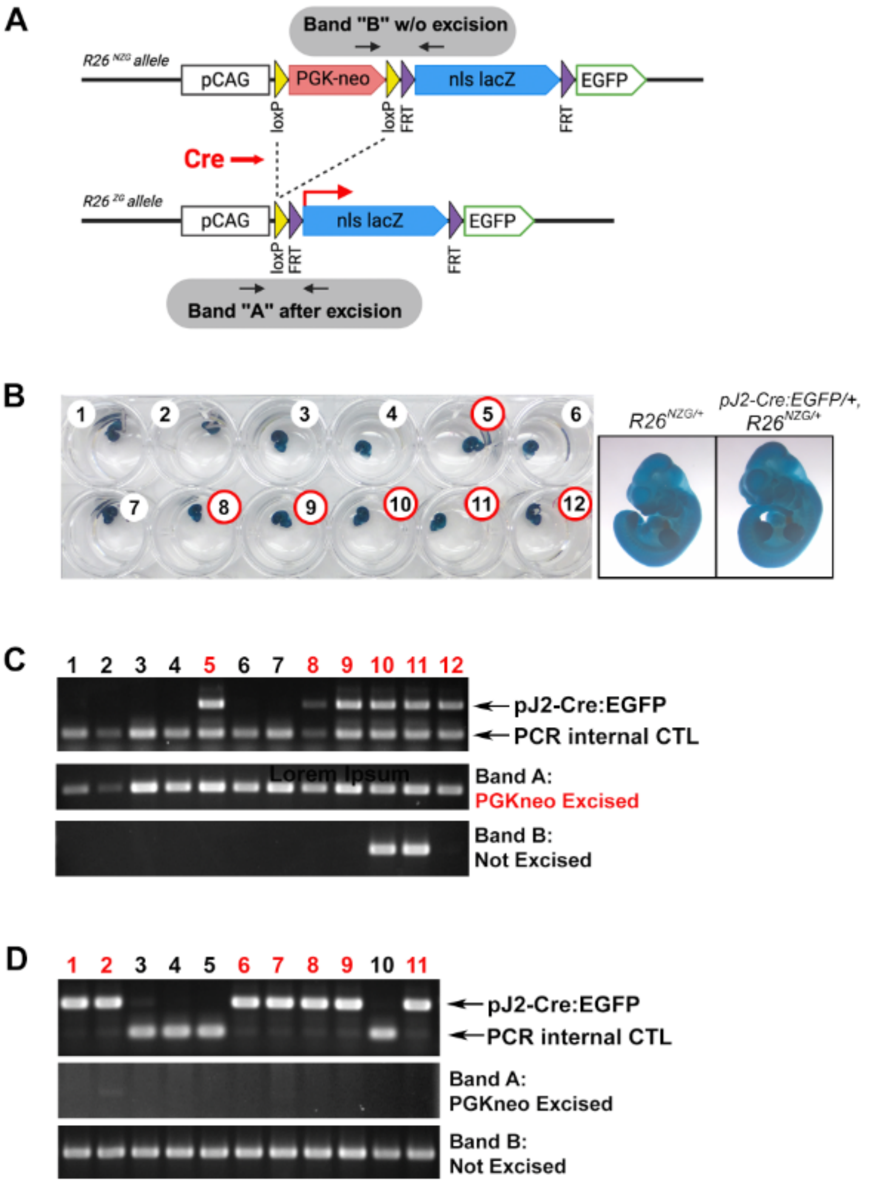

**Figure S1. Female *pJ2-Cre:EGFP/+* mice have maternal Cre expression during oogenesis, leading to excision of the floxed transgene in offspring in the absence of *DUX4* regulatory elements.** A) Schematic of $R26^{NZG}$ transgenes with and without Cre-mediated recombination. Genomic PCR band A shows Cre-mediated excision and genomic PCR band B shows the non-excised transgene. B, C) Heterozygous female *pJ2-Cre:EGFP/+* mice were crossed with homozygous $R26^{NZG}/R26^{NZG}$ males and 12 embryos (E11.5) were analyzed by B) X-gal staining and C) genomic PCR using amniotic sac and umbilical cord genomic DNA. Double transgenic (red numbers) and $R26^{NZG/+}$ heterozygotes (black numbers) all showed Cre-mediated excision (Band A) regardless of inheritance of the pJ2-Cre:EGFP transgene. D) Heterozygous male *pJ2-Cre:EGFP/+* mice were crossed with homozygous female $R26^{NZG}/R26^{NZG}$ mice and 11 embryos (E13.5) were similarly analyzed by genomic PCR. Double transgenic embryos (red numbers) did not show any paternal Cre activity.

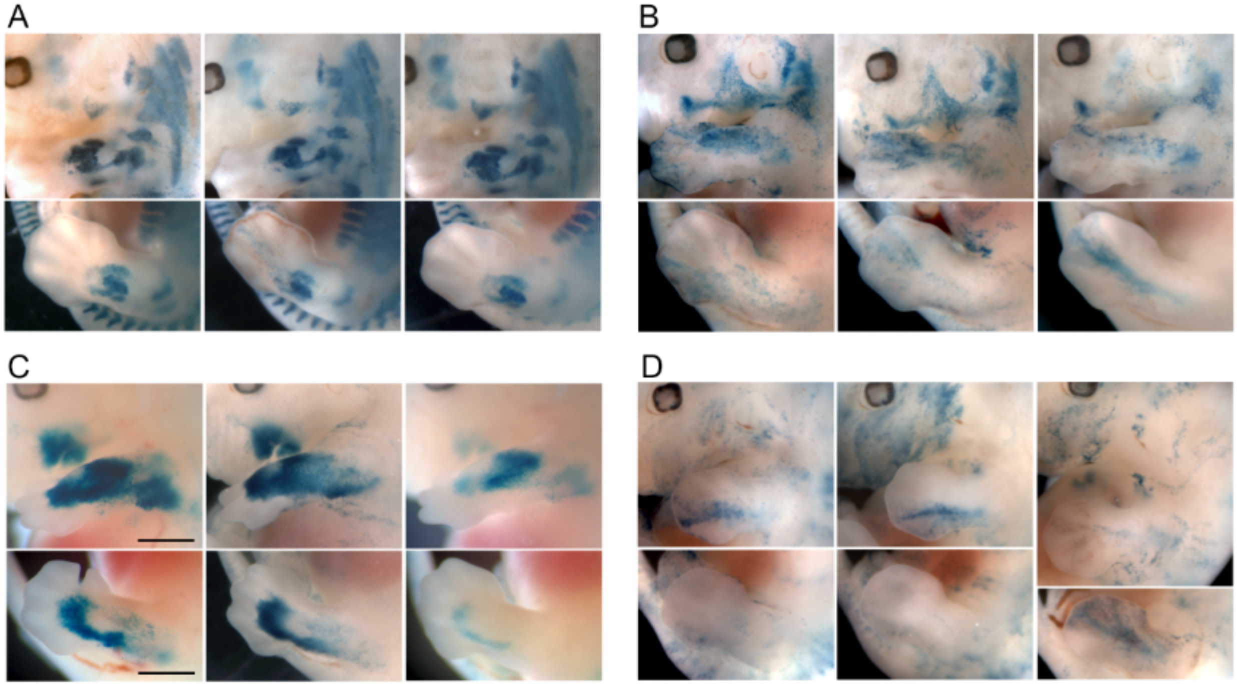

**Figure S2. Embryonic activity of *DUX4* regulatory elements among littermates.** X-gal staining of three representative E13.5 embryos in the same litter of A) *ACTA1-cre/+, R26*$^{NZG/+}$, B) line #6, C) line #7, and D) line #11 *pJ2-Cre:EGFP/+, R26*$^{NZG/+}$. X-gal staining in face and forelimb (top panels), and in hindlimb of the same embryos (bottom panels). Scale bar: 1 mm.

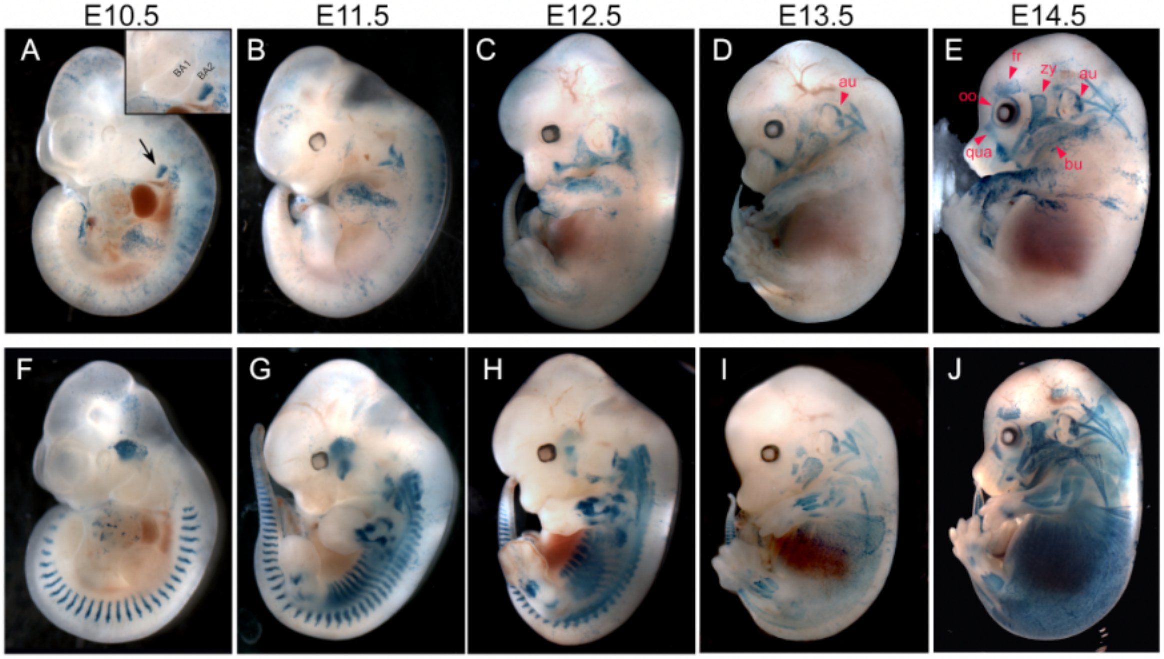

**Figure S3. *DUX4* regulatory elements in line #6 are active in the cell lineage of facial expression muscles during embryonic development.** X-gal staining of line #6 *pJ2-Cre:EGFP/+;* $R26^{NZG/+}$ embryos (A-E), and *ACTA1-cre/+;* $R26^{NZG/+}$ embryos (F-J). A) At E10.5, line 6 shows the X-gal signal in $2^{nd}$ branchial arch (BA2, arrow). Closeup of BA2 signal is shown. B-E) Developmental activity of *DUX4* regulatory elements in line #6. X-gal staining suggests the cell lineage found in BA2 develops to form facial skeletal muscles located closer to the surface. Abbreviation of facial expression muscles in D and E are as follows: au, auricularis; bu, buccinator; fr, frontalis; oo, orbitalis oculi; qua, quadratus labii; zy, zygomaticus.

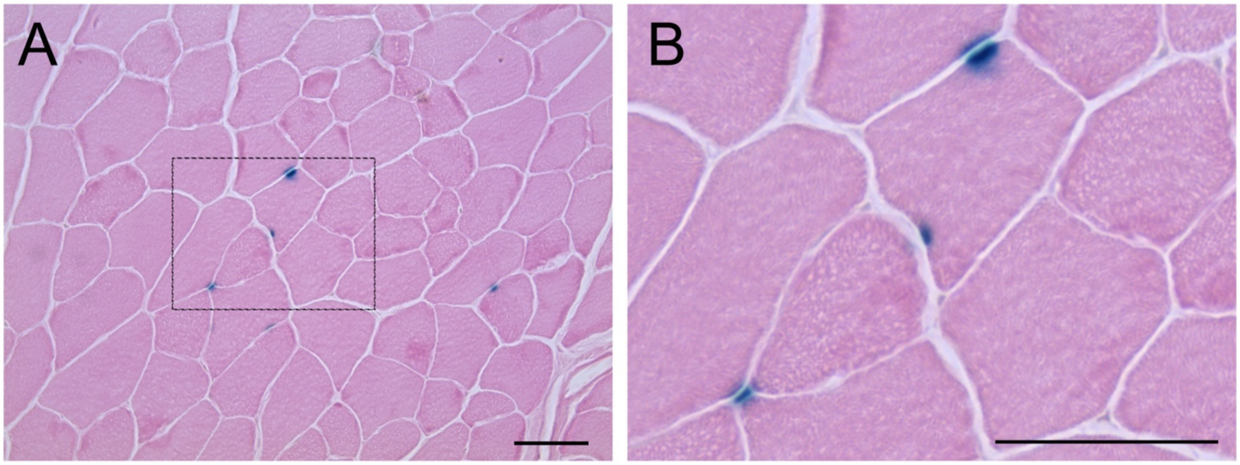

**Figure S4. Activity of *DUX4* regulatory elements in healthy skeletal muscle.** A) X-gal staining of uninjured line #7 *pJ2-Cre:EGFP/+; R26*$^{NZG/+}$ mice at >8 weeks of age shows myonuclear X-gal staining indicating LacZ expression. B) Enlargement of the rectangle in A. Scale bar: 50 µm.

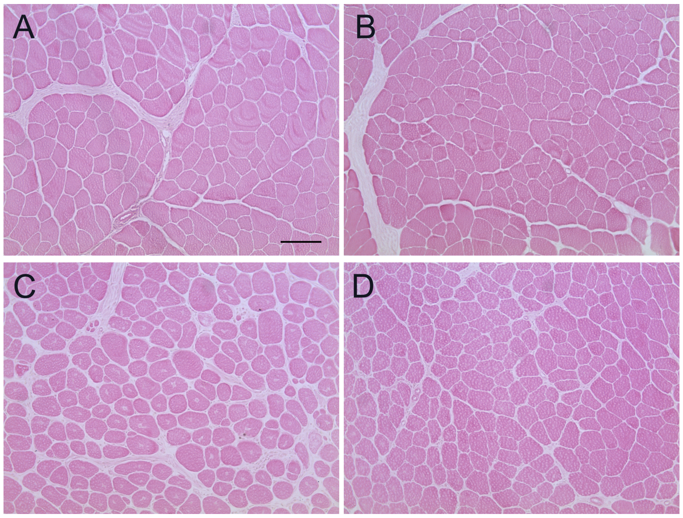

**Figure S5. R26$^{NZG/+}$ negative control for X-gal staining.** X-gal and eosin staining of healthy (A and B) and injured TA muscles (C and D) at 10 days after barium chloride injection. Scale bar = 100 μm.

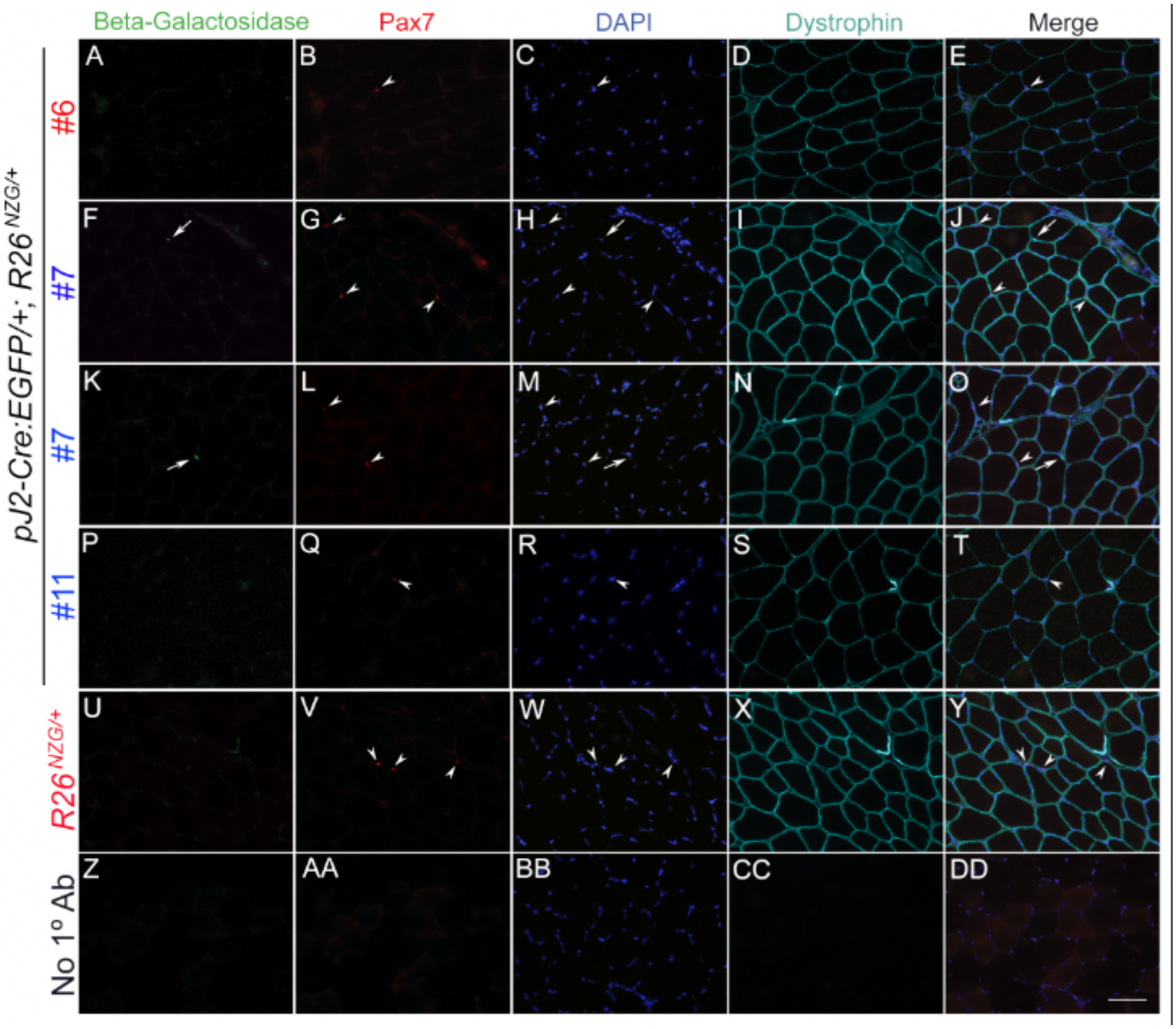

**Figure S6. The interstitially localized cells in which *DUX4* regulatory elements are active are not Pax7+ muscle satellite cells.** TA muscle sections from male (bllue) and female (red) in indicated lines of *pJ2-Cre:EGFP/+; R26*$^{NZG/+}$ and *R26*$^{NZG/+}$ mice were immunostained for beta-galactosidase (green, A, F, K, P, U), Pax7 (red, B, G, L, Q, V), and dystrophin (aqua, D, I, N, S, X), and stained with DAPI (blue, C, H, M, R, W, BB) to show nuclei. Z-DD) no primary antibody control. White arrows show beta-galactosidase-positive nuclei and corresponding nuclei in DAPI panels; white arrowheads show Pax7 positive nuclei and corresponding nuclei in DAPI panel. These are the same immunofluorescent images used in Fig 7 but shown in smaller magnification. Scale bar = 50 μm.

**Table S1.**

Oligonucleotide sequences.

| Name | Sequence (5'-3') |
|---|---|
| Genotyping | |
| R47 #6 5' integration site F | gttggcaacttcagtgcaatc |
| R43 #6 5' integration site R | ccgagtcccggtcttttgtc |
| R44 #6 3' integration site F | cggaaggacatatgggaggg |
| R45 #6 3' integration site R | gtggggtgttgaaatctccc |
| R56 #7 5' integration site F | ggtggtcctggcttatttctc |
| R57 #7 5' integration site R | ggaatgtgtttgtgaagcacc |
| R58 #11 5' integration site F | gagccattgtggtgtttacct |
| R59 #11 5' integration site R | cagaacctgaagatgttcgcg |
| R54 #11 3' integration site F | ctctatgaacttccatgggacc |
| R55 #11 3' integration site R | cagaataccaatagcacaggc |
| Cre F | ttactgaccgtacaccaaaatttgcctgc |
| Cre R | cctggcagcgatcgctattttccatgagtg |
| Mapping analysis | |
| #6 Cre TLA F | ggagtttcaataccggagat |
| #6 Cre TLA R | attacgtatatcctggcagc |
| #7, #11 p13E-11 TLA F | cattcgaactcacaggca |
| #7, #11 p13E-11 TLA R | aactcccagtatctccttca |
| #6, #7, #11 GFP TLA F | caacagccacaacgtctata |
| #6, #7, #11 GFP TLA R | cgtccttgaagaagatggt |
